# Incursion of killer sponge *Terpios hoshinota* on the coral reefs of Lakshadweep archipelago

**DOI:** 10.1101/617662

**Authors:** Rocktim Ramen Das, C.R Sreeraj, Gopi Mohan, K.R Abhilash, Deepak Samuel, Purvaja Ramachandran, Ramesh Ramachandran

## Abstract

Our study documents the outbreak of killer sponge *Terpios hoshinota* on the coral reefs of Lakshadweep archipelago and highlights that the killer sponge has further extended its territory in the isolated atolls of the Indian subcontinent.

## Introduction

Coral killing sponges have the potential to overgrow live corals, eventually killing the coral polyps, and thus leading to an epidemic^1^. The cyanobacteriosponge *Terpios hoshinota* Rützler & Muzik, 1993 first reported from Guam^1^ and later described from the coral reefs of the Ryukyu archipelago (Japan)^2^, is identified by its gray to blackish encrustations. Since its first occurrence, it has been observed in several coral reef localities around the globe viz. the Great Barrier Reef^3^, Papua New Guinea^4^, Taiwan^5^, Philippines^6^, Indonesia^7^, South China Sea^8–10^, Thailand^6^, Palk Bay (PB)/Gulf of Mannar (GOM) (India)^11–13^, Maldives^14^, Mauritius^15^ and our present observation, confirms that the species has further extended its habitat into the pristine atolls of Lakshadweep (Figure 1) (Indian Ocean) and requires urgent attention.

**Figure 1.**
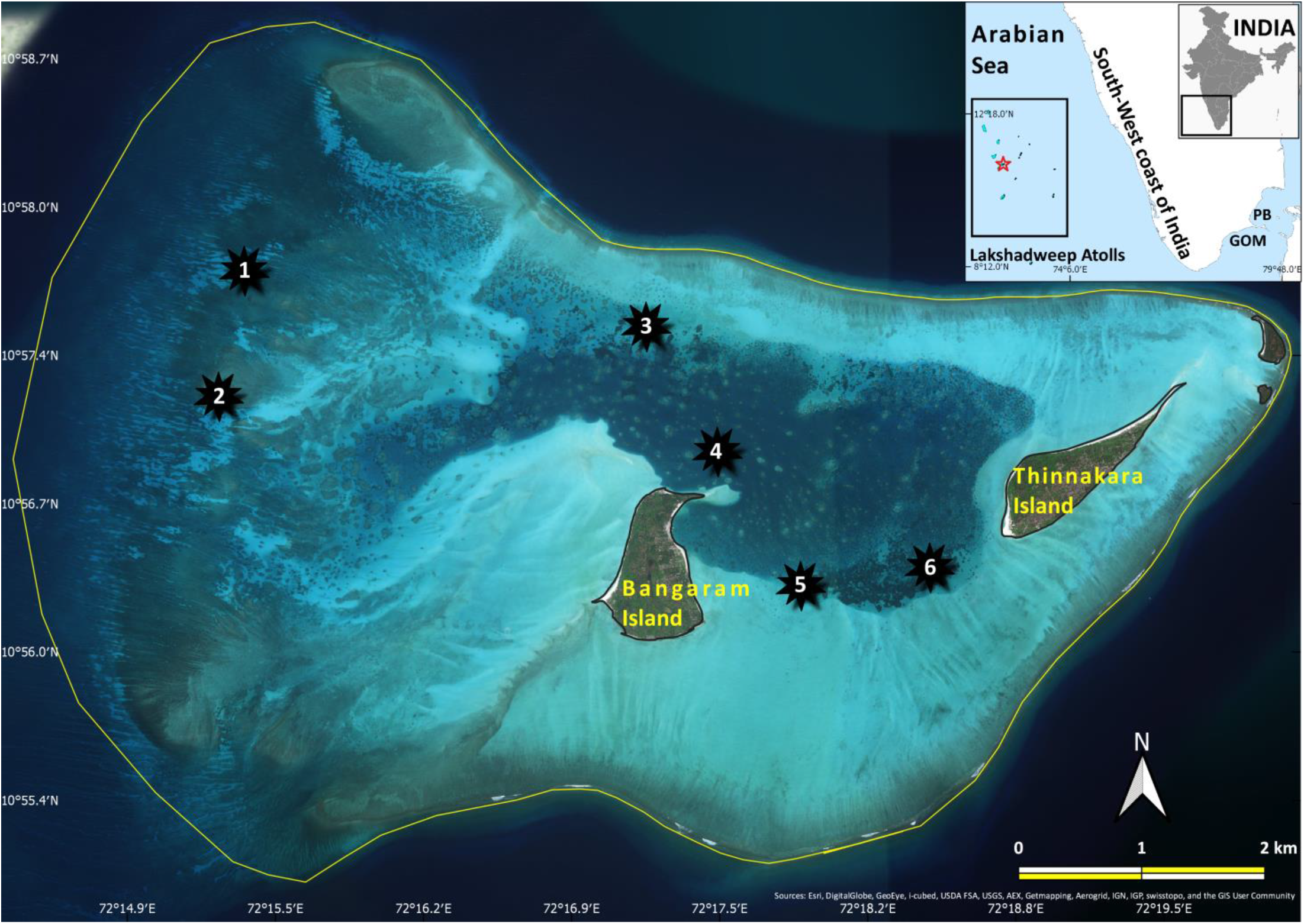
Bangaram & Thinnakara Atoll (Red star); killer sponge infested areas (1-6) (QGIS 3.0.1)

## Results and Discussion

During the coral reef surveys conducted at Lakshadweep in November 2016, *Terpios hoshinota* was observed overgrowing on several colonies of *Acropora formosa, Acropora palifera, Cyphastrea* sp., *Favia lizardensis* and *Porites lutea* (Figure 2 and 3) in the atoll encircling Bangaram and Thinnakara Islands. Out of 33 stations surveyed, 6 stations exhibited the presence of *T.hoshinota* incrustations (Figure 1). The coral colonies in the atoll were patchy and the depth varied between 2 to 12 meters. As depth increased, (i.e. >5 m) large boulder corals were observed whereas the shallow regions (<5 m) had greater coral diversity. Certain areas consisting of large *Acropora* beds, rocks, rubbles and dead reef were also observed. The affected corals displayed grayish/blackish encrustations of *T.hoshinota* forming a mat-like layer on live corals taking the shape of the coral in all cases. The osculum in the sponge, a primary character with a radiating network was clearly visible, the thickness of the mat was less than 1mm (Figure 1a). It was observed that the encrustations were propagating laterally and infecting the other live coral colonies. Further, in some colonies along with *T.hoshinota,* algal presence was also noted (Figure 3a) but the sponge was absent in the colonies which were completely covered with turf (Figure 3b). Other associated communities such as ascidians and clams remain unaffected but interestingly the calcareous serpulid tubes, though overgrew by the *Terpios* like sponge, the animal was unharmed^15^ (Figure 2d). Environmental parameters assessed in the atoll revealed that the area was unpolluted with an optimum level of dissolved oxygen (5.04~8.21 mg/l), and low turbidity (0.3 to 0.8 NTU). Sea surface temperature (SST) during the survey was 28.2°~30.1°C. It is important to note that, Bangaram and Thinnakara is one of the few atolls in Lakshadweep where tourism is permitted, as a result, limited amounts of diving and other water-related recreational activities can be seen in the area.

**Figure 2.**
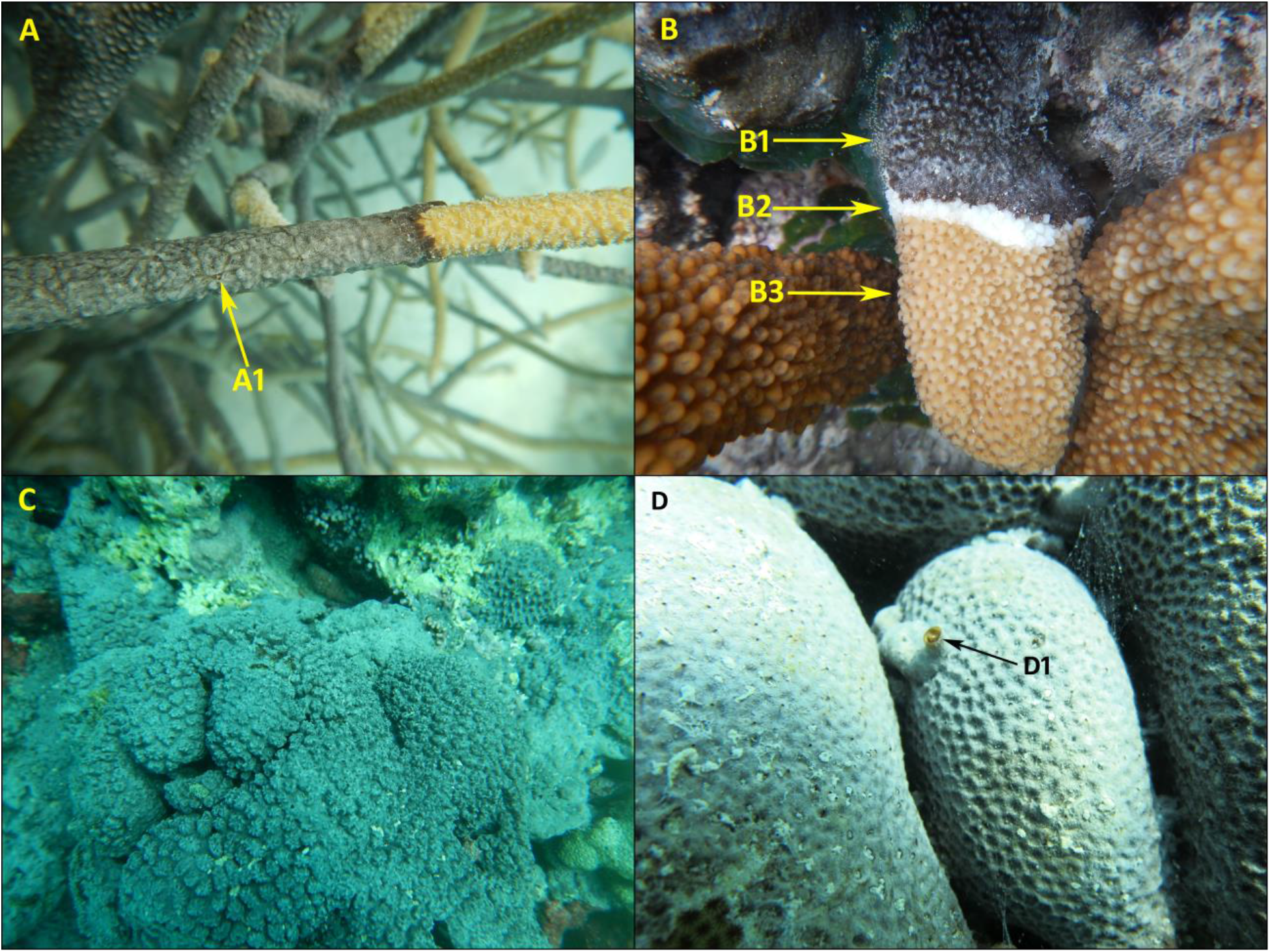
Incrustations of *T.hoshinota* (A) *Acropora formosa,* A1. *T.hoshinota* exhibiting osculum with radiating networks (B) Incrustation on *A.palifera,B1. T.hoshinota* mat, B2. Bleached ring, B3. Live coral (C) *T.hoshinota* taking shape of a Coral *(Cyphastrea* sp.) (D) *Terpios hoshinota* overgrowing calcareous serpulid_tubes, animal unaffected.

**Figure 3.**
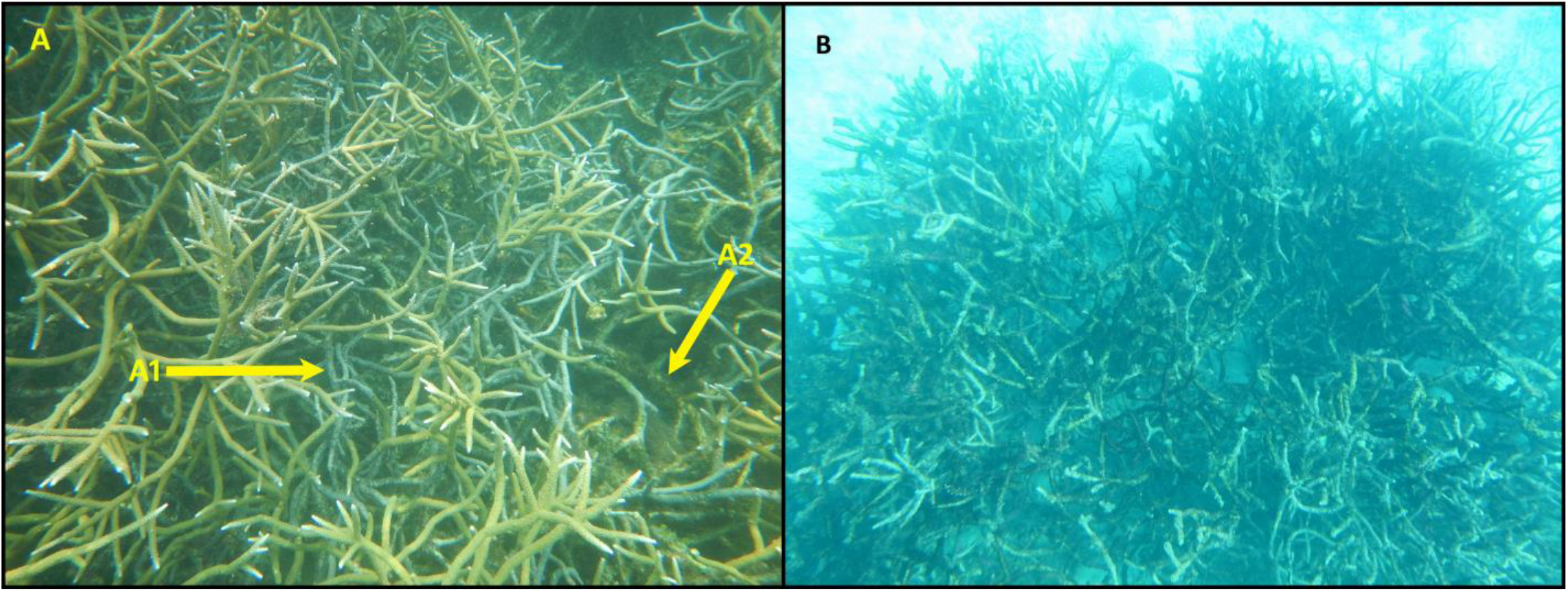
*Acropora* colonies (Station 3) A1. *T.hoshinota* A2. Algae (B) *Acropora* colonies (Station 5) completely over grown by turf algae, killer sponge absent.

Previous studies^2, 11^ suspected that the outbreak of *T.hoshinota* is related to increased water turbidity or due to high anthropogenic stress/pollution and its close proximity to mainland as seen in the south eastern reefs of India (~800km from Lakshadweep)^11–13^, in Guam^2, 6^ and in Green island^17^. However, a similar conclusion cannot be applied in the case of Lakshadweep because of its isolated geography^18^ and with comparatively less anthropogenic activities. As a result, our observation contradicts the above statements and is more in line with the findings of Shi et al.^8^ who observed *T.hoshinota* outbreak in unpolluted areas of Yongxing Island (South China Sea), highlighting the difficulty in establishing negative co-relationship between water quality and killer sponge outbreak (Sun-Yin Yang pers comm.). In terms of host selectivity, the killer sponge has affected several coral species in different parts of the world^1,11,13,15^ and in the reefs of Palk Bay (PB), it has affected all genus surveyed^11^. In Vaan Island (GOM) the dominant genus *Montipora* was the most susceptible^13^. Our observation though could not reveal any specific host coral selectivity, we can speculate that the dense *Acropora* (ACB) beds in station 3, 5 and 6 were more easily overgrown because the killer sponge prefers branching corals as reported from Mauritius^15^. We would further conclude that the coral composition in any specific location may play an important role in determining its host.

*Terpios hoshinota* is a belligerent contender for space^6^ and is known to overgrow corals from its base where it interacts with turf algae^15^. In stations 3, 5 and 6 (Figure 3a), we observed ACB colonies with both algae (e.g. *Ceratodicyton repens?, Dictyota* sp.) and killer sponge, however, the observation of a massive turf algal cover dominating an area of ~0.35km of *Acropora* bed in *T.hoshinota* occurrence site station 5 and 6 indicates towards a much complex ecological scenario. Such complexity between sponges, corals and algae can be only understood through long term monitoring. Gonzalez-Rivero et al.^19^ stated that sponges can act as a potential group that can facilitate and influence coral-algal shifts by acting as a “*third antagonist*” as observed in Glover’s atoll (Belize). Based on our knowledge of the life history of *T.hoshinota* we can just hypothesize the scenario in station 5 and 6 as follows: - (1) *T.hoshinota* invades and overgrows the *Acropora* beds → (2) The coral dies which is followed by the death of the killer sponge → (3) Turf algae takes over. (Figure 3a, b). Moreover, reports of turf algae being a dominant component in the atolls^18^ might indicate towards a faster transition. Globally Elevated SST is a major threat to coral reefs^20^, and the reefs of India^21,22,23^ including the atolls^24^ are no different. With reports indicating that elevated SST has already depleted the coral ecosystem of Lakshadweep, which was evident during 1998^18^, 2010^24^ and 2016^20^ mass bleaching events, it can provide an opportunity for sponges to invade^25^. Dynamics of waterflow^18^ may also play a crucial role in this regard.

## Conclusion

Our findings confirm that the infestation of *T.hoshinota* on the coral colonies of Lakshadweep is currently limited to only Bangaram and Thinnakara atoll as it was not observed in other islands surveyed. There is although a possibility that the killer sponge could invade nearby atolls as seen in other regions^1, 26^. Our observation is indeed the first documentation of *T.hoshinota* on the reefs of Lakshadweep and can be regarded as a baseline for subsequent studies in the islands. Further, to protect the reefs of Lakshadweep, coral health monitoring is required which will allow us to understand the nature of occurrence, distribution, the impact and the causative factors of the killer sponge as well as other coral diseases as per which various coral reef management plans can be initiated.

## Acknowledgements

This study was undertaken as part of the grant-in-aid project of “Mapping of Ecologically Sensitive Areas (ESAs) and Critically Vulnerable Coastal Areas (CVCAs) along the Coast of India”, “[F.No. 22-29/2008-WBICZM-IA-III; 19 June 2014]”. The authors acknowledge the financial and technical support provided by MoEF&CC, Government of India, and the World Bank under the India ICZM Project and are grateful to Dr. Shao-Lun Liu (Tunghai University) for identifying few algae species, to Dr. Sen-Lin Tang (Biodiversity Research Center, Academia Sinica) for providing ref. 10 and to Dr. Sung-Yin Yang (Shimoda Marine Research Center, University of Tsukuba) for ref. 17 and additional information on the outbreak of killer sponge in the South China sea.

